# High-resolution cryo-EM structures of the *E. coli* hemolysin ClyA oligomers

**DOI:** 10.1101/558338

**Authors:** Wei Peng, Marcela de Souza Santos, Yang Li, Diana R. Tomchick, Kim Orth

## Abstract

Pore-forming proteins (PFPs) represent a functionally important protein family, that are found in organisms from viruses to humans. As a major branch of PFPs, bacteria pore-forming toxins (PFTs) permeabilize membranes and usually cause the death of target cells. *E. coli* hemolysin ClyA is the first member with the pore complex structure solved among α-PFTs, employing α-helices as transmembrane elements. ClyA is proposed to form pores composed of various numbers of protomers. With high-resolution cryo-EM structures, we observe that ClyA pore complexes can exist as newly confirmed oligomers of a tridecamer and a tetradecamer, at estimated resolutions of 3.2 Å and 4.3 Å, respectively. The 2.8 Å cryo-EM structure of a dodecamer dramatically improves the existing structural model. Structural analysis indicates that protomers from distinct oligomers resemble each other and neighboring protomers adopt a conserved interaction mode. We also show a stabilized intermediate state of ClyA during the transition process from soluble monomers to pore complexes. Unexpectedly, even without the formation of mature pore complexes, ClyA can permeabilize membranes and allow leakage of particles less than ∼400 Daltons. In addition, we are the first to show that ClyA forms pore complexes in the presence of cholesterol within artificial liposomes. These findings provide new mechanistic insights into the dynamic process of pore assembly for the prototypical α-PFT ClyA.

## Introduction

As the largest family of bacterial toxins, pore-forming toxins (PFTs) represent important factors for bacterial virulence. The protein family is characterized by the pore-forming activity and these types of proteins exist in other kingdoms of life (Dal Peraro and van der Goot, 2016; Iacovache et al., 2010). Proteins exhibiting pore-forming activity, named as pore-forming proteins (PFPs), cause permeation of the target membrane, disruption of cellular homeostasis and death of target cells. With respect to protein structure, PFTs can be divided into two classes, α-PFTs, and β-PFTs, depending on the secondary structure of transmembrane elements of α-helices and β-barrels, respectively (Dal Peraro and van der Goot, 2016; Iacovache et al., 2010).

*E. coli* cytolysin A (ClyA), also known as hemolysin (HlyE) or silent hemolysin A (SheA), belongs to the α-PFT class (Dal Peraro and van der Goot, 2016; Mueller et al., 2009). ClyA has been shown to be responsible for the hemolytic activity of certain strains of *E. coli*, including the pathogenic strain *E. coli* O157 (del Castillo et al., 1997; Oscarsson et al., 1996; Ralph et al., 1998). Homologs of ClyA are also found in other pathogenic bacteria, including *Salmonella typhi* and *Shigella flexneri* (del Castillo et al., 1997; Oscarsson et al., 2002). Distinct from many secreted toxins and effectors, ClyA is observed to be delivered by outer-membrane vesicles (OMVs) (Wai et al., 2003). As the first α-PFT with the structure of a pore complex determined, ClyA represents the prototype in the ClyA α-PFT subfamily (Mueller et al., 2009). Other subfamily members include NheA and Hbl-B from *Bacillus cereus*, and Cry6Aa from Bacillus thuringiensis, which share obvious sequence or structural similarities with ClyA (Dementiev et al., 2016; Ganash et al., 2013; Madegowda et al., 2008). ClyA is a 34 kDa soluble monomer, containing mainly α-helices with only one β-tongue that transforms into α-helical elements in the pore complexes (Mueller et al., 2009; Wallace et al., 2000).

Various low-resolution EM analyses suggest that the ClyA pore complexes form 6-, 8- or 13-fold symmetrical structures (Eifler et al., 2006; Tzokov et al., 2006). However, the crystal structure of ClyA pore complex displays only a 12-meric composition (Mueller et al., 2009). Due to the limitations of the protein crystal lattice, which self-selects for a single oligomeric species, it is reasonable to propose the existence of ClyA pore complexes of different oligomeric states. Here we report the cryo-EM structures of ClyA pore complexes in the presence of n-Dodecyl-β-D-Maltoside (DDM) as dodecamer, tridecamer, and tetradecamer at the resolution of 2.8 Å, 3.2 Å and 4.3 Å, respectively. These structures clearly reveal the diverse organizations of ClyA pore complexes, with high-resolution electron microscopy density maps contributing to more accurate assignments of amino acid residues for the dodecamer. Structure comparisons indicate a rigidity of protomers when assembled into pore complexes. We also show that a mild detergent, digitonin, is able to facilitate the transition of ClyA from soluble monomer to an intermediate state. With the presence of cholesterol within artificial liposomes, ClyA forms pore complexes. Overall, our studies reveal the pore assembly mechanism involving intermediate oligomers for the formation of a prototypical α-PFT of ClyA with varying numbers of protomers.

## Results

### Assembly of the ClyA Pore Complexes

Detergents, including DDM and n-Octyl-β-D-Glucopyranoside (β-OG), have been shown to induce the *in vitro* formation of the ClyA pore complexes (Eifler et al., 2006; Mueller et al., 2009; Tzokov et al., 2006). We first tested the effect of DDM on the assembly of the ClyA complexes. As shown in **Figure S1A**, DDM caused a dramatic shift of the ClyA protein elution peak to a smaller elution volume with size exclusion chromatography (SEC), indicating the formation of large oligomers of ClyA pore complexes. Characteristic ClyA pore complexes were observed with negative staining EM images (**Figure S1B**). Interestingly, the ClyA pore complexes from an earlier or later eluted fraction with SEC exhibited slightly bigger or smaller central pore size, respectively, implying a heterogeneous composition of ClyA pore complexes. The variance of ClyA pore size was also observed in cryo-EM images (**Figure S4A**). Soluble ClyA monomer was previously found to assemble into a 13-meric pore complex in the presence of DDM (Eifler et al., 2006), but a dodecamer was observed in the crystal structure (Mueller et al., 2009). EM images of ClyA pore complexes from a β-OG/lipid mixture displayed 6- or 8-fold symmetry (Tzokov et al., 2006). To determine the oligomerization states of the complexes we obtained, we took advantage of cryo-EM for further investigation.

### Structure Determination of the ClyA Pore Complexes

We pooled all fractions containing ClyA pore complexes in DDM after SEC for cryo-EM grid preparation. With this sample numbered as sample #1 (**Figure S1C**), we were able to perform grid screening and collect micrographs on a Talos Arctica operated at 200 kV. After data processing, we successfully obtained characteristic 2D class averages, with a relatively small dataset of 410 micrographs containing 22,998 auto-picked particles (**Figure S2A** and **S2C**). Two top view images (or bottom view, as they cannot be differentiated) clearly indicated the presence of both dodecamer and tridecamer (**Figure S2A**). 3D reconstruction with further classified particles produced low-resolution density maps of the dodecamer and the tridecamer, at resolutions of 6.8 Å and 7.5 Å, respectively. Based on the particle numbers of the dodecamer and the tridecamer, the majority of the complexes were 12-meric, representing approximately 80% of all particles, while only 11% were 13-meric. These observations confirm the existence of 13-metric ClyA pore complexes (Eifler et al., 2006), albeit at a low concentration in this preparation.

Digitonin is generally considered to be a mild detergent and has been used successfully for cryo-EM structure determinations of membrane proteins (Gong et al., 2016; Peng et al., 2016; Qian et al., 2017). Therefore, we replaced DDM with digitonin after pore formation and added an additional step of Ni^2+^-NTA affinity purification before SEC (sample #2, **Figure S1C**). The data from sample #2 obtained on the same microscope (Talos Arctica) yielded a dodecamer density map at the resolution of 3.9 Å from fewer particles (**Figure S2D**). The tridecamer was lost during this sample preparation process, as evidenced by the lack of 2D class averages for a tridecamer (**Figure S2B** and **S2D**). In the hopes to visualize the tridecamer, we subsequently adopted a purification protocol by skipping the Ni^2+^-NTA affinity purification and directly applying the DDM complex to SEC with digitonin in the buffer (sample #3, **Figure S1C**). Data for sample #3 were collected on a Titan Krios (300 kV) to obtain micrographs of better quality.

With the data from sample #3, we were able to observe not only the tridecamer in addition to the dodecamer but also a new oligomer with 14-fold symmetry (**Figure 1A**, **S3**, and **S4B**). Final 3D reconstruction resulted in EM density maps of the dodecamer, the tridecamer and the tetradecamer at the resolution of 2.8 Å, 3.2 Å and 4.3 Å, respectively (**Figure 1B**, **S3**, and **S4**; **Table S1**). The three density maps clearly exhibit distinct secondary structural elements, demonstrating that DDM is capable of inducing the formation of ClyA pore complexes of multiple oligomeric states. The near-atomic resolution maps of the dodecamer and the tridecamer allowed reasonable model building with accurate side chain assignments. Although the tetradecamer map was resolved at a relatively low resolution, the model was still built based on the dodecamer structure model with identical protomer fold and inspection of the electron microscopy density maps for assignment of bulky residues (**Figure 1B** and **S5**; **Table S1**).

**Figure 1.**
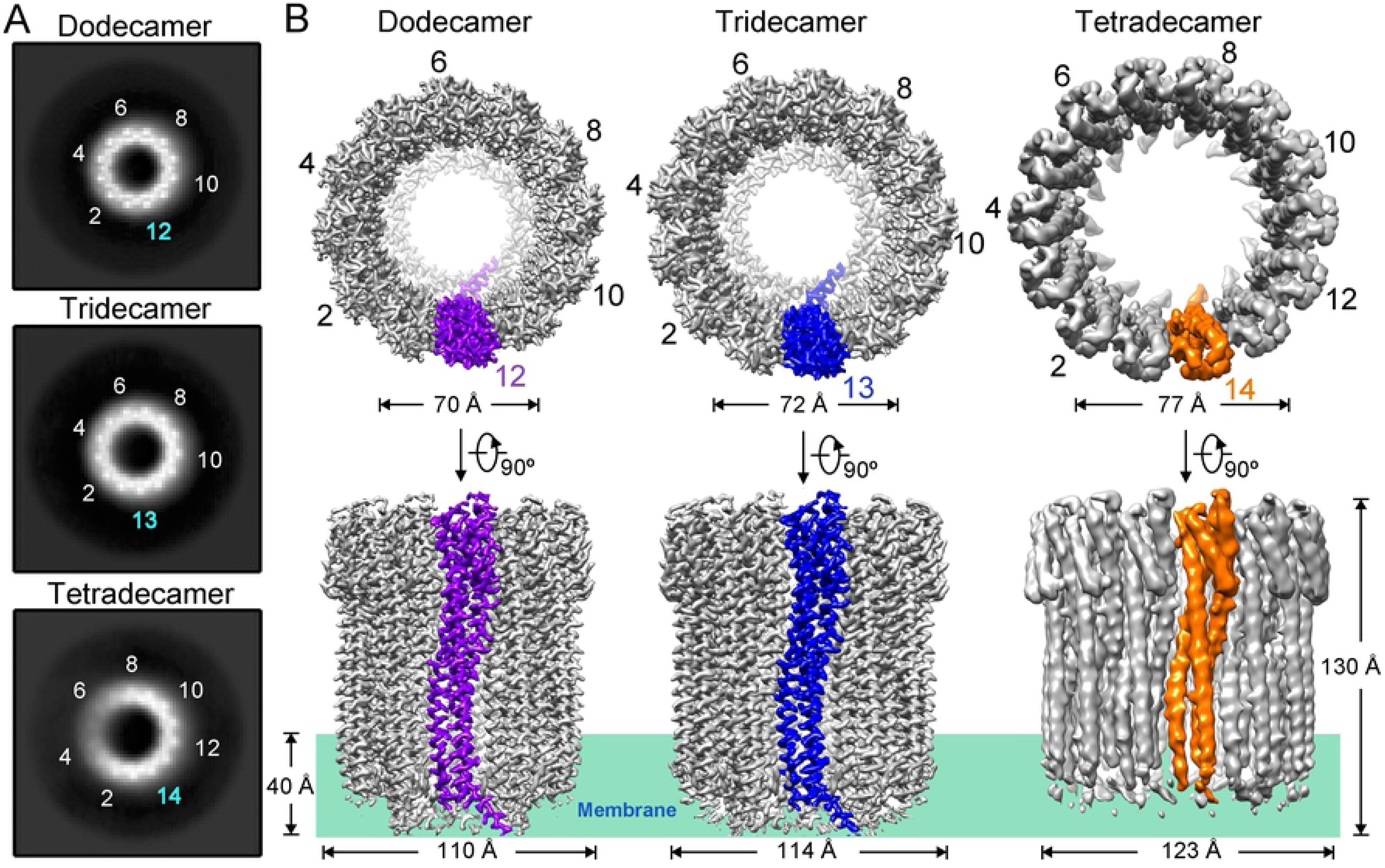
Cryo-EM structures of the ClyA pore complexes. (**A**) Top view (or bottom view) of the dodecamer, the tridecamer and the tetradecamer from 2D class averages of the ClyA pore complexes. The numbers of symmetric features are indicated. (**B**) The final three-dimensional reconstruction electron microscopy density maps of the dodecamer, the tridecamer, and the tetradecamer at the resolution of 2.8 Å, 3.2 Å and 4.3 Å. Top views and side views are shown. One protomer from each oligomer is shown in purple, blue or orange for clarity.

### Structure Comparison of the ClyA Pore Complexes

For membrane proteins, the transmembrane regions are usually more stable, and the local resolutions are better compared to those of the soluble regions. However, this is not the case for the ClyA pore complexes, since the local resolutions of all three transmembrane regions are worse than that of the soluble regions (**Figure S4E**). This may result from the incomplete transmembrane helices from αC and αF, with the turning point located within rather than outside of the membrane, making the transmembrane regions unstable (**Figure 2A**, black arrow). Although the N-terminus of αA1 helix from the tetradecamer is even more disordered and not well resolved, we built the model similar to the dodecamer (**Figure 1B**, **S4E**, and **S5**).

**Figure 2.**
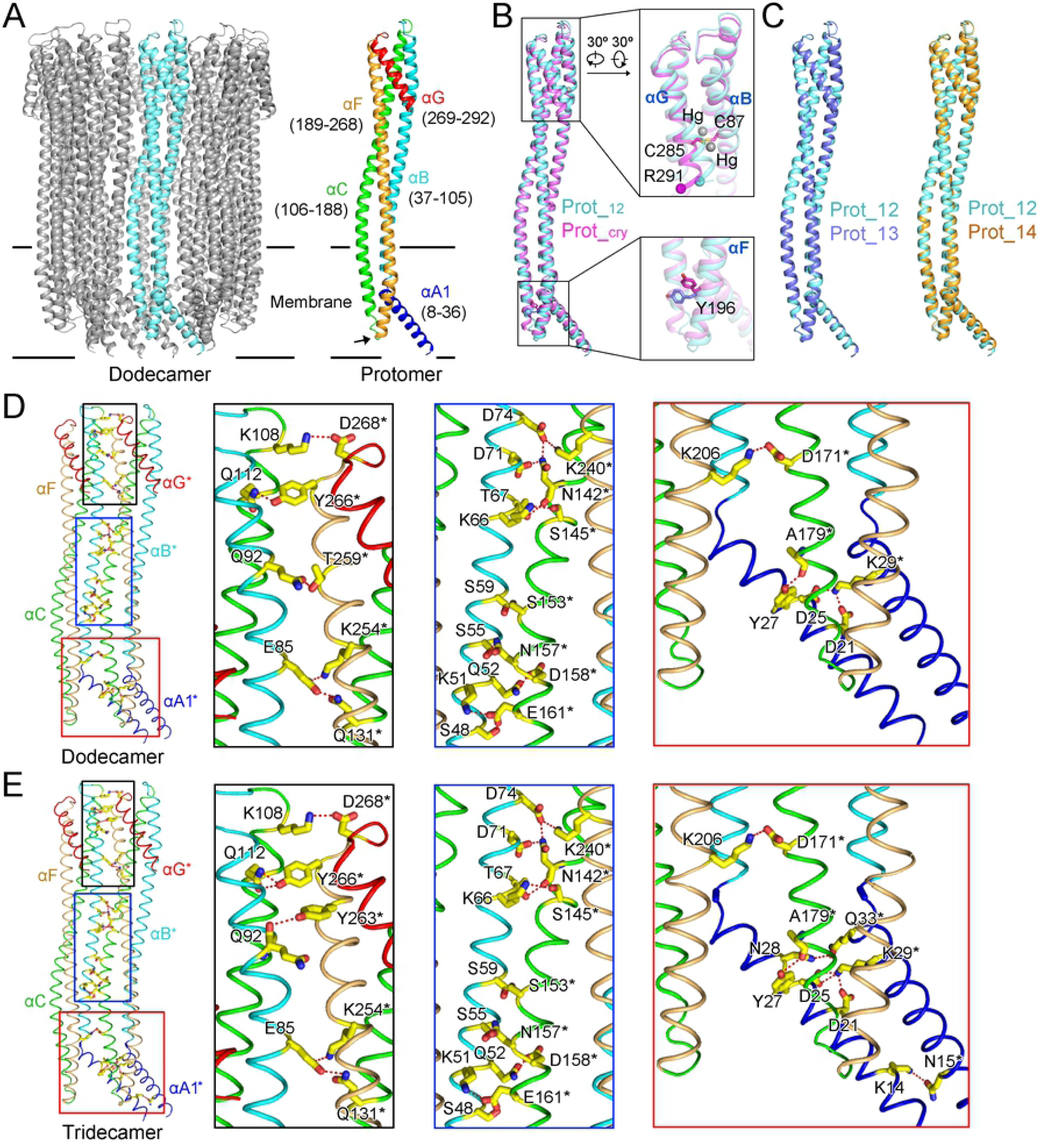
Structure comparison of the ClyA dodecamer, tridecamer, and tetradecamer complexes. (**A**) Illustration of secondary elements within one protomer from the dodecamer. Five α-helices (αA1, αB, αC, αF, and αG) within one protomer (left) are colored in blue, cyan, green, orange and red (right). The corresponding amino acid boundaries are indicated, with the short loops divided and assigned to the linked helices for simplicity. The turning point of αC and αF is indicated by an arrow. The same coloring paradigm is applied in **Figure 2D**, **2E**, and **S7**. (**B**) Comparison of one protomer from the dodecamer (Prot_12, in cyan) with that from the crystal structure (Prot_cry, in magenta). The PDB accession code for the crystal structure is 2WCD. The two mercury atoms in the crystal structure are shown as grey balls. Cα atoms of R291 are indicated as balls. (**C**) Comparison of Prot_12 with a protomer from the tridecamer (Prot_13, in blue) and the tetradecamer (Prot_14, in orange). (**D**, **E**) Extensive hydrogen bonds and salt bridges mediate the interaction between neighboring protomers in the dodecamer and the tridecamer. Residues involved are labeled and shown as sticks. Residues from different protomers are distinguished with or without “*”.

The height of the dodecamer, the tridecamer or the tetradecamer is approximately 130 Å. The outer diameter varies from 110 Å to 123 Å as ClyA pore complexes incorporate more protomers. The inner diameters near the extracellular edge of the pore are 70, 72 and 77 Å for the dodecamer, the tridecamer, and the tetradecamer, respectively. The narrowest part along the pore is located at the membrane, where the inner diameters vary from 40 Å to 52 Å (**Figure 1B** and **S6**).

Each protomer in the ClyA pore complexes is composed of five α-helices, namely αA1, αB, αC, αF, and αG (**Figure 2A**). The crystal structure of the ClyA pore complex was previously solved as a dodecamer at 3.3 Å (Mueller et al., 2009). The EM density map of the dodecamer presented here is at 2.8 Å, representing an improved density and structure model (**Figure S5**). One typical example is the side chain of Y196 from αF, which is buried in the membrane. With the cryo-EM density map, we were able to re-assign the side chain in parallel to, rather than pointing toward the hydrophobic membrane environment (**Figure 2B** and **S5**). Sodium ethylmercurithiosalicylate was used as the heavy atom source for structure determination of the crystal structure, and two mercury atoms are bound to the only two cysteine residues; C87 from αB and C285 from αG (Mueller et al., 2009). One protomer from the dodecamer (Prot_12) was superimposed to a protomer from the crystal structure (Prot_cry) (**Figure 2B**). Based on the comparison, it is obvious that the binding of mercury atoms pushes away the C-terminus of the αG helix with a movement distance of 4.6 Å at the Cα of R291 (**Figure 2B**). This represents an artificial distortion of the local structure by heavy atoms, although the overall fold is not affected (Mueller et al., 2009).

Comparison between Prot_12 and a protomer from the tridecamer (Prot_13) or the tetradecamer (Prot_14) shows that the overall fold of the protomer does not change when forming various oligomers, indicating a rigid body protomer assembly mechanism for the ClyA pore complexes (**Figure 2C**). Extensive hydrogen bonds and salt bridges, spanning from the transmembrane regions to the distal soluble regions, mediate the interaction between neighboring protomers within the dodecamer, which is also observed in the crystal structure (**Figure 2D** and **S7**) (Mueller et al., 2009). The interaction paradigm remains highly conserved in the tridecamer, suggesting a single protomer-protomer interaction mode in the ClyA pore complexes (**Figure 2E**).

### An Intermediate State of ClyA during Transition into Pore Complexes

As discussed above, the detergent digitonin was helpful for determining the high-resolution cryo-EM structures of ClyA pore complexes. However, DDM was still necessary for the formation of mature pore complexes before its replacement with digitonin, as incubation of soluble ClyA monomers with digitonin did not promote the formation of ClyA pore complexes (**Figure 3A** and **3B**). Unexpectedly, ClyA incubated with digitonin formed trimer-like complexes (**Figure 3B**). Native PAGE gel confirmed the medium size of the trimer-like complex between the soluble monomer and the mature pore complex (**Figure 3C**). The digitonin-induced trimer probably represents an intermediate state of ClyA during the transition from soluble monomers to mature pore complexes. The fluorescence emission spectra of the intermediate state differ from the soluble monomer and pore complex, suggesting a different conformation and further supporting that the digitonin induced complex represents an intermediate.

**Figure 3.**
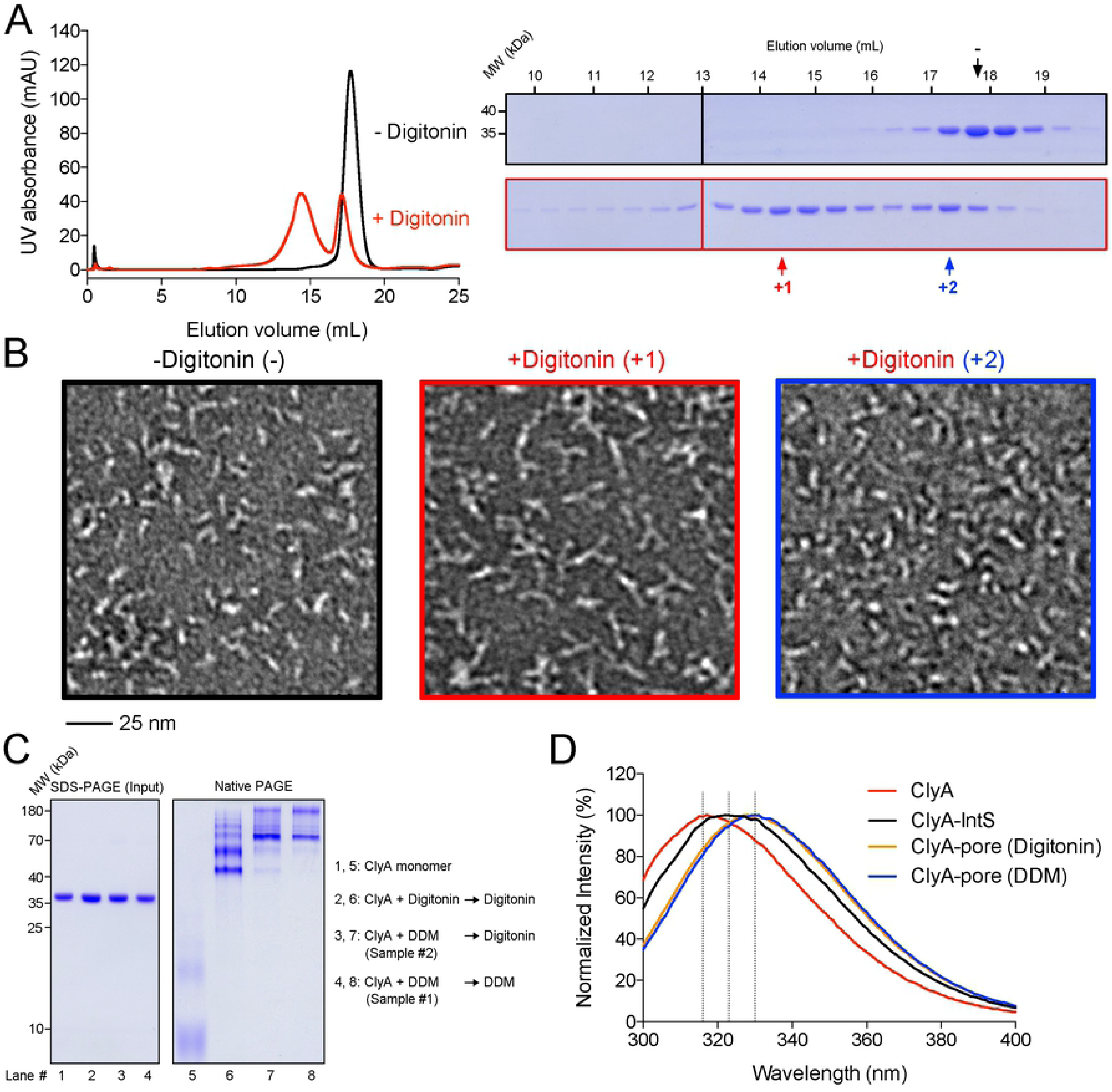
Characterization of a ClyA intermediate state. (**A**) SEC profile of ClyA in the soluble state and an intermediate state. Digitonin (1%) was incubated with ClyA overnight before gel filtration to obtain the intermediate state sample. The elution volumes for the two states were 17.7 mL and 14.4 mL (Superose 6 10/300 GL, GE Healthcare), respectively. (**B**) Negative staining images of ClyA in the form of soluble state and intermediate state. Fractions indicated in (**A**) were applied to negative staining (“-” indicates the soluble protein, “+1” or “+2” indicates the sample with digitonin). (**C**) Native PAGE analysis of ClyA in the soluble state (lane 1, 5), the intermediate state (lane 2, 6), and the state of mature pore complex (lane 3, 4, 7, 8). (**D**) Fluorescence emission spectra of the ClyA monomer, the intermediate state (IntS) and the pore complexes. Wavelengths of 316, 323, and 330 nm are indicated for clear visualization of peak shift.

To further characterize the ClyA trimer, we also collected cryo-EM micrographs on a Talos Arctica. More than 15,000 particles were picked from the images. After data processing, several 2D class averages confirmed the trimeric feature of the intermediate state, though others showed tetrameric feature (**Figure S8A**). 3D classification and reconstruction with approximately 5,000 classified particles resulted in an EM density map at the resolution of 16 Å (**Figure S8B**). With this low-resolution map, details on the protomer structure and the interaction between protomers could not be discerned. However, based on the overall long-rod shape of the ClyA monomer and the protomer (**Figure S8B** and **S8C**), the trimer organization manner must be either head-to-head or tail-to-tail.

The transition of ClyA from soluble monomers to pore complexes involves significant conformational changes of αA1, αA2, αD, β-tongue, and αE, which insert into the membrane, while αB, αC, αF and αG function as the scaffold (**Figure 2A** and **S8C**) (Mueller et al., 2009; Wallace et al., 2000). Partial scaffold elements could self-assemble into ring-like structures without membrane-buried elements (Tzokov et al., 2006), explained by the extensive protomer-protomer interactions (**Figure 2D** and **2E**). Therefore, the corresponding transmembrane elements in the soluble monomer prior to complete rearrangement in the protomer inhibit the assembly of ClyA ring-like pore complex. The intermediate state shown here appears as a trimer, indicating that digitonin is not able to completely disrupt the interactions stabilizing ClyA as a monomer and facilitate the maturation of the ClyA pore complexes.

### The Interaction between ClyA and Liposomes

ClyA is capable of forming pores on artificial liposomes made of porcine brain total lipid extract (BTLE) (Eifler et al., 2006; Tzokov et al., 2006; Wallace et al., 2000). Therefore, we carried out a liposome leakage assay for testing the interaction of ClyA with membrane lipids and its ability to permeabilize liposomes. Liposomes were prepared with 1-palmitoyl-2-oleoyl-glycero-3-phosphocholine (POPC) and 1,2-dioleoyl-sn-glycero-3-phospho-L-serine (DOPS) or BTLE and then tested for the leakage activity of ClyA. A *Vibrio parahaemolyticus* type III secretion system effector, VopQ (known to induce liposome leakage), and its chaperone VP1682, were tested as positive and negative controls, respectively, in the liposome leakage assay (Akeda et al., 2009; Sreelatha et al., 2013). At pH 5.5, VopQ causes fast and dramatic 5(6)-Carboxyfluorescein (CF) dye release from POPC-DOPS liposomes, while at pH 7.4, the leakage activity is less efficient (**Figure S9A**), consistent with previous observations (Sreelatha et al., 2013). Addition of ClyA showed similar leakage activity with POPC-DOPS liposomes (**Figure S9A**), with higher leakage activity at lower pH. The observations may be explained by changes in the protein overall charge, as the estimated pIs of VopQ and ClyA are 5.7 and 5.1, respectively (Sreelatha et al., 2013). Experiments done at a low pH caused the pore-forming proteins to be more positively charged, enabling them to bind more efficiently to the negatively charged liposomes. ClyA also caused leakage of BTLE liposomes, which contained more diverse mammalian cell lipids (**Figure S9B**). Pre-formed ClyA pore complexes were also tested and found to show no obvious leakage activity (**Figure S9B**). This is consistent with observations that mature ClyA pore complexes show little hemolytic activity on erythrocytes (Fahie et al., 2013; Hunt et al., 2008).

We examined whether ClyA formed visible pore complexes on the same liposomes used in our leakage assay by negative staining. As shown in **Figure S9C**, ClyA did not form obvious pore structures on POPC-DOPS liposomes, even though we observed dramatic leakage activity (**Figure S9A**). Since the protein tested contained an N-terminal affinity purification tag (21 amino acids), which would relocate from the outside of membrane to the inside and may impede pore complex formation, we removed the tag and tested ClyA (−tag). In the presence of DDM, the tag-free ClyA protein formed pores similar to those formed by the tagged protein, as expected, based on the fluorescence emission spectra (**Figure S8D**). Even at a higher concentration (5 μM), ClyA (−tag) still could not form pores on POPC-DOPS liposomes (**Figure 4A**). ClyA was previously shown to form pores on BTLE liposomes (Eifler et al., 2006; Tzokov et al., 2006; Wallace et al., 2000). We then tested BTLE liposomes with different lipid composition and did find pore-like structures (**Figure 4A**). Cholesterol has been revealed to facilitate membrane binding and pore complex stabilization of ClyA (Oscarsson et al., 1999; Sathyanarayana et al., 2018). Cholesterol is not listed as a component on the manufacturer’s data sheet, but approximately 60% of BTLE ingredients are unknown. As a common animal membrane component, cholesterol would be expected to be present in the BTLE, explaining the observation that ClyA pores were found on BTLE liposomes. To test more carefully the effect of cholesterol, we prepared POPC-DOPS-cholesterol liposomes and indeed observed pore-like structures on these liposomes (**Figure 4A**).

**Figure 4.**
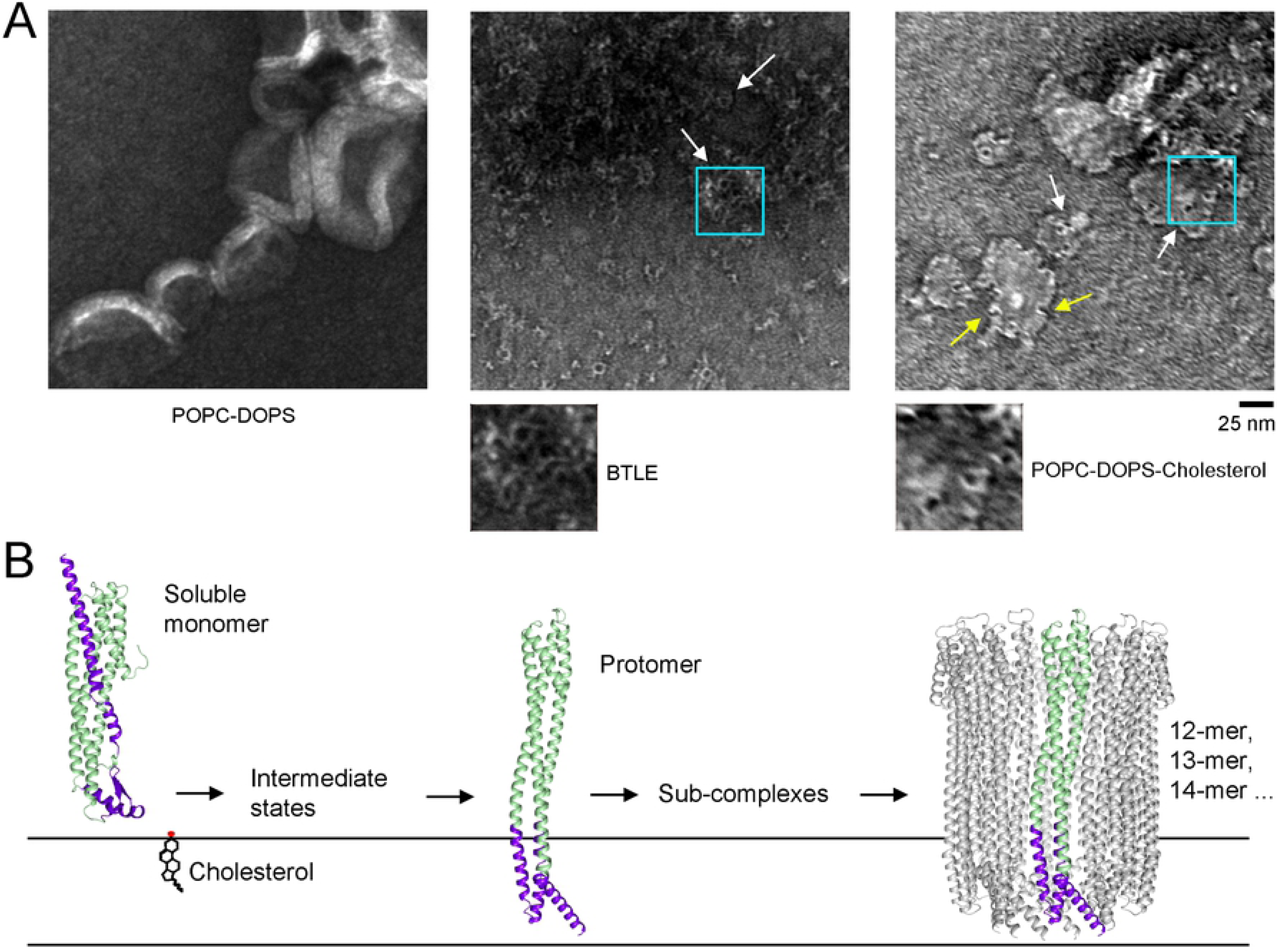
ClyA forms pores on cholesterol-containing liposomes. (**A**) POPC-DOPS, POPC-DOPS-cholesterol or BTLE liposomes were incubated ON with 5 μM ClyA (−tag) at 4 °C (pH 7.4) and examined with negative staining. ClyA forms pore structures on POPC-DOPS-cholesterol and BTLE liposomes, but not on POPC-DOPS liposomes. Representative pores are indicated with white arrows. Regions within the cyan boxes are enlarged for clear visualization. Crescent structures are indicated with yellow arrows. (**B**) Model for the assembly of ClyA pore complexes from the soluble monomer. The corresponding transmembrane elements in the soluble monomer and one protomer are colored purple.

The above observations on the liposome leakage activity and the pore formation on POPC-DOPS liposomes suggest that the leakage activity of a ClyA monomer is not equivalent to the membrane permeability mediated by mature pore complexes of ClyA. Therefore, the leakage activity on liposomes can be uncoupled from the pore-forming activity for ClyA, and possibly for any given protein with pore-forming activity. Moreover, the irrelevance of the leakage activity to the pore-forming activity also indicates that ClyA pore complexes are not essential for leakage of molecules smaller than the 376 Dalton CF dye. The initial interaction between ClyA monomer and membrane would thus cause membrane instability and leakage, even before the formation of mature ClyA pore complexes.

## Discussion

PFTs are important bacterial virulence factors with pore-forming activity on target membranes (Dal Peraro and van der Goot, 2016; Iacovache et al., 2010). As a prototypical α-PFT, ClyA from *E. coli* assembles into pore structures on the membrane with a pore size of 40 Å (Mueller et al., 2009; Wallace et al., 2000). There has been controversy about the organization of ClyA pore complexes with low-resolution EM analysis (Eifler et al., 2006; Mueller et al., 2009; Tzokov et al., 2006). In addition to the dodecamer which has been characterized by protein crystallography (Mueller et al., 2009), here we report high-resolution cryo-EM structures of the tridecamer and the tetradecamer from a single dataset. The density maps allow accurate structural model building and elucidation of (Schubert et al., 2018) the profile of ClyA pore complexes. Based on the cryo-EM structures, we eliminated artifacts such as the movement of the αG helix due to mercury atoms bound in the crystal structure (Mueller et al., 2009). The pore complex protomers from the tridecamer and the tetradecamer exhibit the same fold as observed with the dodecamer, indicating the rigidity of ClyA protomer when assembled into pore complexes. The protomer-protomer interaction paradigm in the dodecamer applies also to the tridecamer, and probably the tetradecamer.

Various oligomerization states for PFTs or PFPs assembly have also recently been observed via cryo-EM single particle analysis, for a two-component α-PFT YaxAB from *Yersinia enterocolitica*, or XaxAB from *Xenorhabdus nematophila*, and a pyroptotic β-PFP of gasdermin A3 (Brauning et al., 2018; Ruan et al., 2018; Schubert et al., 2018). This illustrates the advantage of cryo-EM in determining distinct organization states of PFP complexes, which could not be achieved with protein crystallography (Mueller et al., 2009). The distinct organization states for PFPs are not surprising due to the flexibility or plasticity of high-order assembly.

Outer membrane vesicles (OMVs) are used for export, delivery and, activation of ClyA *in vivo* (Wai et al., 2003). However, ClyA pores on OMVs seemed to exhibit an inverted tail-outside orientation. This orientation is particularly intriguing because it requires shuttling the large ClyA tail – rather than the comparatively small head group – across the vesicular membrane (Wai et al., 2003). The tail-outside orientation of ClyA was observed on BTLE liposomes, which is to be expected, as the ClyA protein was added from outside (data not shown) (Eifler et al., 2006; Tzokov et al., 2006; Wallace et al., 2000). Unlike on BTLE liposomes, ClyA monomer could not assemble into pore complexes on liposomes prepared with *E. coli* total lipid extract (Fahie et al., 2013). An oligomeric pre-pore state may be escorted by OMVs, representing a possible mechanism for ClyA delivery (Fahie et al., 2013). In vitro, ClyA is shown to form assembly-competent protomer, elongate as sub-complexes and, finally, assemble as mature pore complexes (Benke et al., 2015). As a support for this mechanism, crescent sub-complexes are observed on POPC-DOPS-cholesterol liposomes (**Figure 4A**). Thus, the *in vivo* delivery and assembly mechanism of ClyA still requires further investigation.

Treatment with the mild detergent digitonin is able to induce the formation of a putative intermediate state of ClyA and stabilize the state for long time range study. The cryo-EM analysis results imply that the trimer interaction interface is mediated by the heads or the tails. However, due to the low resolution, detailed structural information about the intermediate state could not be addressed yet. Future investigation on intermediate states may benefit from more detergent options and other biophysical methods, as seen with FRET (Benke et al., 2015). Both mature pores and ring-like pre-pores with immature transmembrane elements (non-membrane-inserted) have been observed for β-PFPs, e.g., α-hemolysin and bi-component γ-hemolysin from *Staphylococcus aureus* and gasdermin A3 (Ruan et al., 2018; Song et al., 1996; Yamashita et al., 2014; Yamashita et al., 2011). However, no pre-pore structure has been identified for α-PFPs, even though a high-order oligomer is observed for YaxAB and multiple states are revealed for FraC from *Actinia fragacea* (Brauning et al., 2018; Tanaka et al., 2015). Further investigations on the pre-pore structures of α-PFPs are necessary for the complete illustration of α-PFP assembly mechanism.

ClyA is capable of inducing leakage of liposomes with a simple composition of POPC and DOPS. Cholesterol has been revealed to facilitate membrane binding and pore complex stabilization (Oscarsson et al., 1999; Sathyanarayana et al., 2018), which may explain the pore formation of ClyA on BTLE liposomes but not POPC-DOPS liposomes. By separating the liposome leakage activity from the pore-forming activity, we demonstrate that binding of ClyA is sufficient to cause liposome leakage of small molecules without the formation of mature pores on liposome membranes. This may also be applicable to other PFPs, adding caution to the interpretation of liposome leakage assay data.

Overall, we have demonstrated that ClyA can form pores of various sizes, including a dodecamer, tridecamer, and tetradecamer. We show an intermediate state of ClyA that is induced and stabilized by digitonin. The ClyA monomer, before assembling into pore complexes, can permeabilize membranes and allow leakage of particles less than ∼400 Daltons. Cholesterol facilitates the assembly of the ClyA pore complexes on membranes. These findings provide new insights into the assembly mechanism for ClyA pore complexes (**Figure 4B**).

## Materials and Methods

Detailed material information and protocols are described in the Supplemental Experimental Procedures. Data availability: The atomic coordinates of the dodecamer, the tridecamer and the tetradecamer of ClyA pore complexes have been deposited in the Protein Data Bank with accession codes 6MRT, 6MRU, and 6MRW, respectively. The corresponding electron microscopy density maps have been deposited in the Electron Microscopy Data Bank with accession codes EMD-9212, EMD-9213, and EMD-9214, respectively.

## Author Contributions

Conceptualization, W.P. and K.O.; Investigation, W.P., M.S.S., Y.L.; Writing – Original Draft, W.P.; Writing – Review & Editing, W.P., M.S.S., Y.L., D.R.T. and K.O.; Resources, D.R.T. and K.O.; Funding Acquisition, K.O.; Supervision, K.O.

## Acknowledgments

We thank Dr. Xiaochen Bai and the Cryo-Electron Microscopy Facility (CEMF), under management by Dr. Daniela Nicastro and Dr. Daniel Stoddard, for technical assistance and support for cryo-EM data collection. We thank the Electron Microscopy Core Facility (EMCF) for assistance with negative staining grid preparation and image obtainment. We thank the Structural Biology Laboratory (SBL) for assistance with cryo-EM grid preparation and screening. We thank BioHPC for providing computational resources for cryo-EM image processing. We thank the Hooper lab for assistance with liposome leakage assay and fluorescence spectroscopy. We thank Dr. Chad Brautigam for insightful discussions and members of the Orth lab for insightful discussions and editing. This work was funded by the Welch Foundation grant I-1561 (KO); Once Upon a Time…Foundation (KO); National Institutes of Health Grant R01 GM115188 (K.O). K.O. is a W.W. Caruth, Jr. Biomedical Scholar with an Earl A. Forsythe Chair in Biomedical Science. D.R.T. is an Effie Marie Scholar.

